# SEQURNA enhances FLASH-seq gene detection while eliminating DTT dependence

**DOI:** 10.64898/2025.12.12.693841

**Authors:** Irina Khven, Mariana Ribeiro, Sara Crausaz, Simone Picelli

**Affiliations:** Institute of Molecular and Clinical Ophthalmology Basel, Basel, Switzerland; Department of Ophthalmology, University of Basel, Basel, Switzerland

**Keywords:** SEQURNA, RNase Inhibitor, FLASH-seq, scRNA-seq, benchmarking

## Abstract

Effective RNase inhibition is critical for single-cell RNA-sequencing, yet commercial recombinant RNase inhibitors (RRIs) require reducing agents for stability and impose substantial costs. Here, we systematically benchmark SEQURNA, a synthetic thermostable RNase inhibitor, against commercial alternatives using FLASH-seq in human retinal organoids and peripheral blood mononuclear cells (PBMCs). SEQURNA at 0.5–1 U/μl achieved 14–50% higher gene detection than Takara RRI, with the greatest improvement in low-RNA PBMCs. Surprisingly, DTT supplementation at standard concentrations (5–10 mM) significantly impaired gene detection across all SEQURNA concentrations without improving RNA quality metrics, challenging established reverse transcription protocols. SEQURNA preserved biological heterogeneity, maintained sample stability during one-month storage at −80°C, and reduced reagent costs by 70%. We recommend SEQURNA at 1 U/μl without DTT as an optimized formulation that simultaneously enhances data quality and cost-effectiveness for full-length single-cell RNA sequencing.

## Background

Single-cell RNA sequencing (scRNA-seq) has transformed our ability to dissect cellular heterogeneity in complex tissues, yet RNA preservation remains a critical bottleneck that fundamentally limits data quality [1]. RNA degradation during cell collection, storage, and library preparation introduces technical artifacts manifesting as reduced gene detection, elevated mitochondrial transcript content, and spurious mapping patterns [2,3]. The small copy numbers of individual transcripts in single cells render scRNA-seq particularly vulnerable to degradation-induced losses, making effective RNase inhibition essential for accurate transcriptome capture.

Commercial recombinant RNase inhibitors (RRIs) have served as the standard solution, yet they present significant limitations [4]. These protein-based inhibitors require reducing agents such as dithiothreitol (DTT) at 5–10 mM concentrations to maintain their leucine-rich repeat structure and prevent oxidation of critical cysteine residues [5,6]. Furthermore, RRIs exhibit restricted specificity, primarily targeting pancreatic-type RNases with femtomolar affinity but remaining ineffective against RNase T1 and other non-mammalian nucleases. While SUPERase• In offers broader spectrum inhibition, it remains protein-based and shares the same thermolability constraints [7,8]. These limitations, combined with substantial costs that contribute to reagent expenses 10–20 times higher than bulk RNA experiments, present barriers to broader scRNA-seq adoption particularly in resource-limited settings [9].

SEQURNA, a synthetic thermostable RNase inhibitor composed of non-toxic organic molecules, offers a promising alternative by eliminating DTT dependence and maintaining activity through heat cycles up to 72°C [10]. The original study demonstrated successful substitution of RRIs in Smart-seq protocols with maintained or enhanced library quality. However, systematic evaluation across cell types with varying RNA content and optimization of concentration parameters in diverse single-cell methods was not performed. Here, we benchmark SEQURNA against commercial alternatives using FLASH-seq [11] in human retinal organoids and human peripheral blood mononuclear cells (PBMCs), representing cells with moderate-to-high and low RNA content, respectively.

## Results and Discussion

We tested SEQURNA at six concentrations (0.2, 0.5, 0.75, 1, 1.25, and 1.5 U/μl in the final RT-PCR reaction, hereafter indicated only as “U”) alongside commercial alternatives (Takara RRI, Takara oxidation-resistant RRI, Watchmaker Genomics RRI) and no-inhibitor controls (Supplementary Figure 1A). To assess storage stability, retinal organoid cells were processed either immediately after sorting (fresh) or following one-month storage at −80°C in lysis buffer containing the respective RNase inhibitors. After quality filtering and computational downsampling to uniform depth, we retained 1,040 retinal organoid cells and 796 PBMCs for downstream analysis.

Gene detection patterns revealed distinct SEQURNA concentration optima depending on cell type and expression threshold (Figure 1A–D). In retinal organoids, lower SEQURNA concentrations (0.2–1 U) achieved maximal breadth of transcriptome coverage, with 0.2 U and U detecting 5,066 ± 1,102 and 4,908 ± 1,221 genes per cell, respectively (mean ± SD, >0 read threshold), comparable to or exceeding Takara RRI (4,322 ± 1,065 genes). Notably, 1 U SEQURNA maintained robust performance with 4,634 ± 1,076 genes while demonstrating superior mapping quality (68.3% uniquely mapped reads, Figure 1E). Higher concentrations showed reduced total gene detection (1.5 U: 3,744 ± 858 genes). However, when examining robustly expressed genes (>10 reads threshold), mid-to-high concentrations (0.75-1.25 U)demonstrated superior performance, suggesting these concentrations reduce low-level background while preserving biologically relevant transcripts (Supplementary figure 2A). The no-RRI control exhibited the poorest performance across all thresholds (3,615 ± 1,140 genes), confirming the critical importance of RNase inhibition.

**Figure 1.**
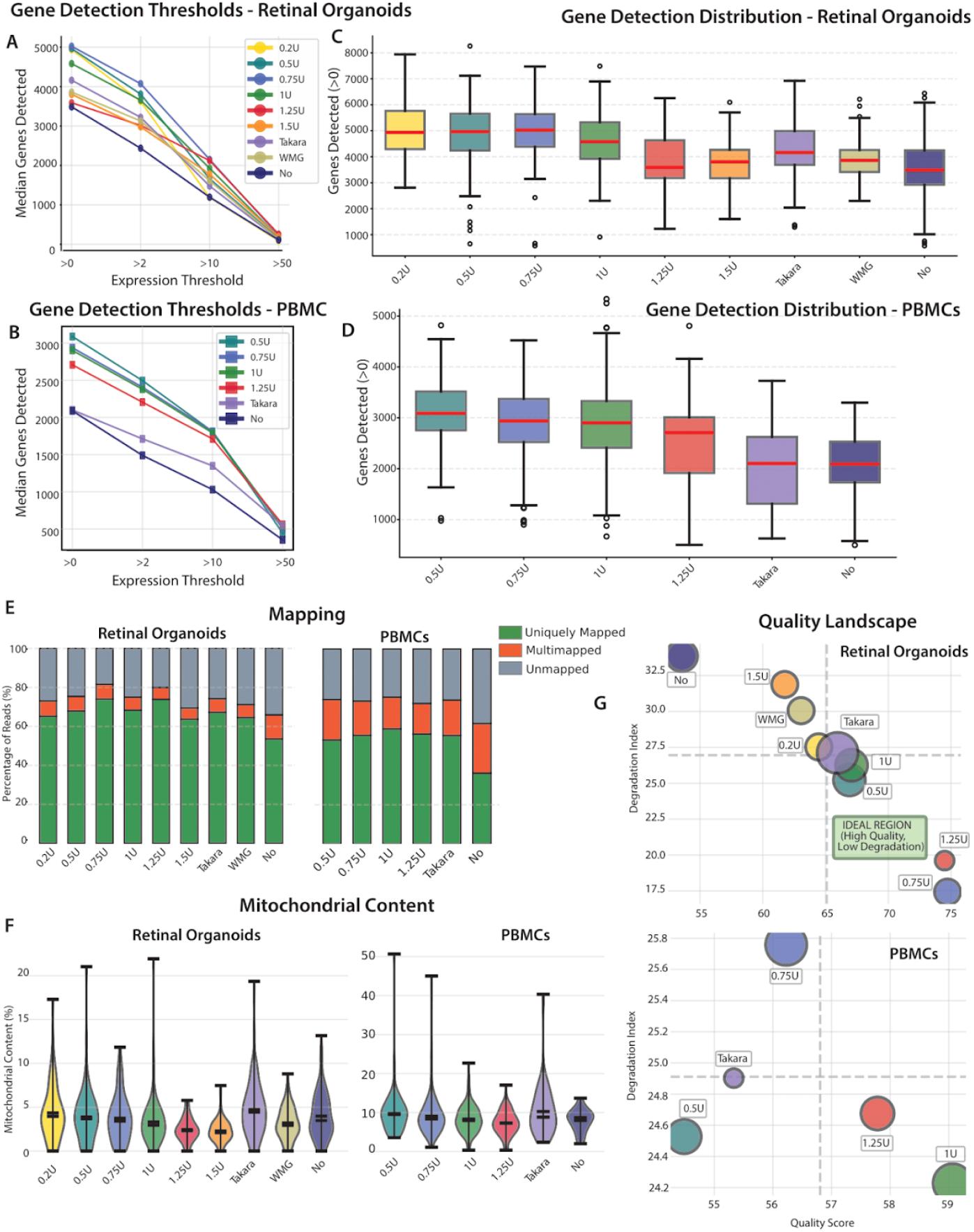
SEQURNA performance across cell types and concentrations. **(A, B)** Median genes detected at increasing expression thresholds (>0, >2, >10, >50 reads) for retinal organoids (A) and PBMCs (B) across SEQURNA concentrations (0.2–1.5 U), commercial RRIs (Takara, WMG), and no-inhibitor controls. **(C, D)** Distribution of genes detected per cell (>0 read threshold) in retinal organoids (C) and PBMCs (D). Boxplots show median, interquartile range, and outliers. **(E)** Mapping efficiency showing percentage of uniquely mapped, multimapped, and unmapped reads for each condition in retinal organoids (left) and PBMCs (right). **(F)** Mitochondrial transcript content (% of total reads) across conditions. Violin plots show distribution with median indicated. **(G)** Quality landscape plotting composite quality score against degradation index for retinal organoids (top) and PBMCs (bottom). Circle size represents number of cells; dashed lines and shaded region indicate ideal performance zone (high quality, low degradation). All samples were downsampled to uniform sequencing depth prior to analysis. WMG, Watchmaker Genomics RRI; No, no-inhibitor control.

In PBMCs, which present a more challenging model due to lower RNA content and predominantly quiescent transcriptional state, SEQURNA at 0.5 – 1 U yielded optimal results. SEQURNA at 0.5 U detected 3,095 ± 636 genes per cell, while 1 U achieved 2,876 ± 831 genes with notably improved mapping efficiency (58.9% uniquely mapped) and the highest splice junction detection among all conditions (9,672 high-confidence junctions per cell). Both substantially outperformed Takara RRI (2,061 ± 825 genes) and no-RRI controls (2,041 ± 717 genes). The performance advantage was more pronounced in PBMCs than retinal organoids, with 0.5 U SEQURNA achieving 50% higher gene detection than Takara RRI compared to 14% improvement in retinal organoids, highlighting SEQURNA’s particular value for challenging low-RNA samples.

Mapping efficiency analysis revealed substantial differences between conditions and cell types (Figure 1E). In retinal organoids, mid-range SEQURNA concentrations achieved the highest unique mapping rates, with 0.75 U and 1.25 U reaching 74.0% and 73.9% respectively, significantly outperforming Takara RRI (67.3%) and the no-RRI control (53.6%). Multimapping rates were consistently low across SEQURNA conditions (5.8–7.9%), whereas no-RRI controls showed elevated multimapping (12.4%), indicative of degradation-induced ambiguous read mapping. In PBMCs, 1 U SEQURNA achieved the highest unique mapping (58.9%), dramatically exceeding the no-RRI control (36.2% unique, 25.6% multimapped), demonstrating effective RNase inhibition is critical in low-RNA samples.

Splice junction detection revealed that PBMCs exhibited 1.7-2.0-fold higher junction counts than retinal organoids across all SEQURNA conditions (Supplementary figure 2C). In PBMCs, 1 U SEQURNA detected 9,672 high-confidence junctions per cell, substantially exceeding Takara RRI (7,045) and no-RRI controls (3,734). The ratio of annotated to novel junctions remained favorable across all conditions (>95% annotated in retinal organoids, >98% in PBMCs), indicating high data quality with minimal spurious splicing artifacts (Supplementary figure 1H).

RNA quality metrics remained favorable across all SEQURNA conditions (Figure 1F-G). Mitochondrial transcript content was low in retinal organoids (median 2.28–4.34%), with mid-range concentrations showing the lowest contamination. At 1 U SEQURNA, mitochondrial content was 3.31%, slightly lower than Takara RRI (4.73%). Ribosomal RNA content was uniformly low across all conditions (Supplementary Figure 1G). Composite quality scores derived from alignment statistics confirmed superior SEQURNA performance, with mid-range concentrations achieving optimal scores (67–74 in retinal organoids) substantially exceeding no-RRI controls (52). Cold storage for one month at −80°C did not significantly compromise data quality (Supplementary figure 1D-F), with comparable recovery rates between fresh (93.2%) and stored samples (95.7%), supporting batched processing workflows.

We systematically evaluated DTT at 0, 5, and 10 mM across multiple SEQURNA concentrations (0.5, 0.75, 1, 1.25 U). Control conditions were tested at 5 mM DTT following manufacturer recommendations. This analysis revealed a consistent negative effect of DTT on gene detection, most pronounced in retinal organoids (Figure 2A–D). In retinal organoids, DTT had a statistically significant dose-dependent effect across all SEQURNA concentrations tested. At 0.5 U SEQURNA, gene detection decreased from 5,468 genes (0 mM DTT) to 4,964 genes (5 mM) and 4,494 genes (10 mM; p<0.01). This pattern was consistent across all concentrations tested (Figure 2C). Notably, mitochondrial transcript content decreased significantly with DTT addition in retinal organoids (Figure 2B), dropping from ∼5% at 0 mM to ∼2–3% at 10 mM DTT. However, this modest reduction does not compensate for the substantial loss in gene detection. In PBMCs, the negative effects of DTT on gene detection were less consistent across conditions (Figure 2D), though still evident at 1U SEQURNA concentration.

**Figure 2.**
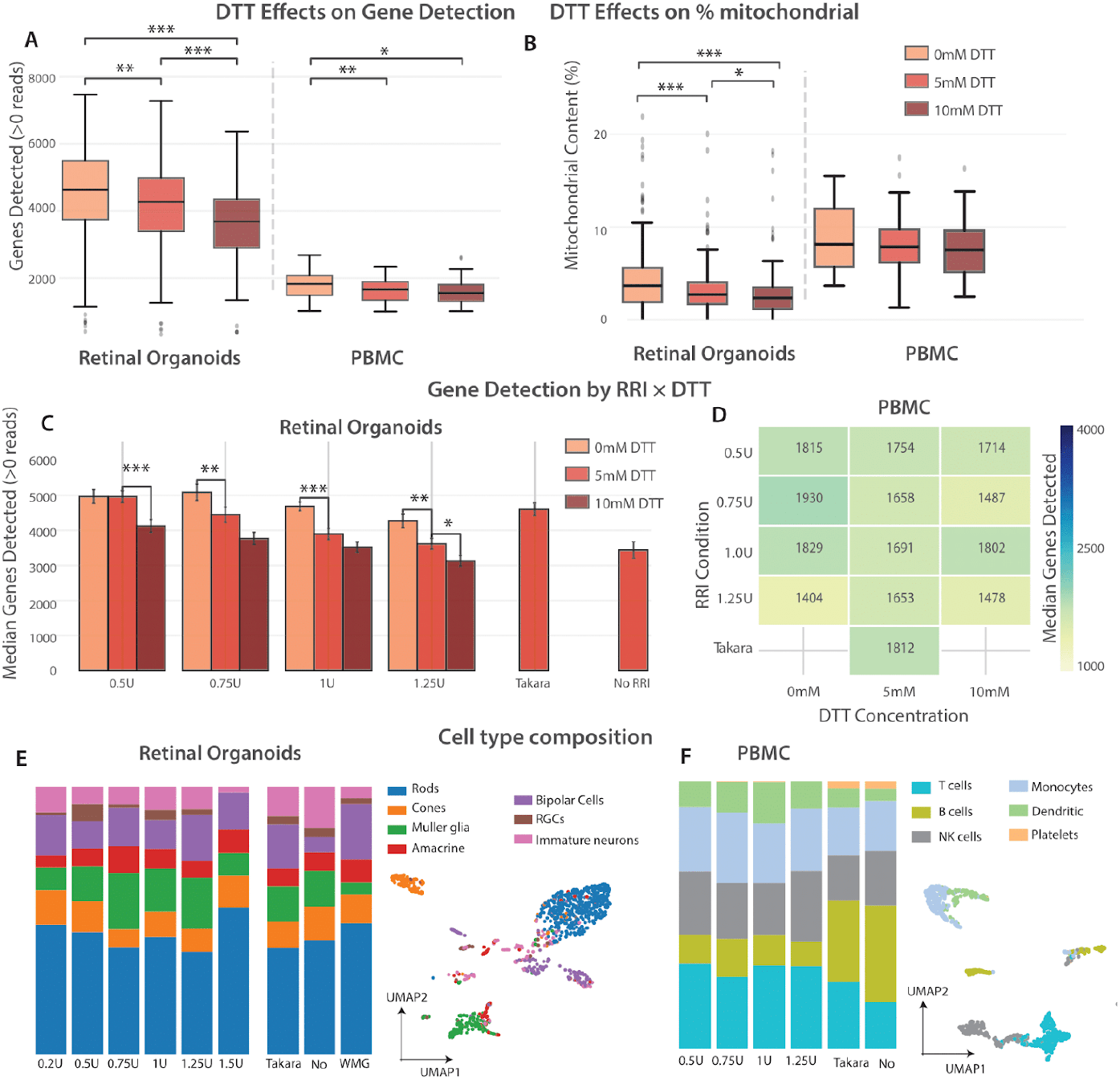
DTT supplementation impairs gene detection without improving RNA quality, while cell type composition is preserved across conditions. **(A)** Gene detection (>0 read threshold) across DTT concentrations (0, 5, 10 mM) in retinal organoids and PBMCs. **(B)** Mitochondrial transcript content (%) across DTT concentrations. Boxplots show median, interquartile range, and outliers. **(C)** Median genes detected per SEQURNA concentration (0.5–1.25 U) and controls (Takara, no RRI) at each DTT concentration in retinal organoids. Statistical comparisons indicate differences between DTT conditions within each RRI group. **(D)** Heatmap of median genes detected in PBMCs across RRI and DTT conditions. Takara RRI was tested only at 5 mM DTT per manufacturer recommendations. **(E, F)** Cell type composition across conditions for retinal organoids (E) and PBMCs (F). Stacked bar charts show proportions of annotated cell types; UMAP projections display expected separation of cells by cell type. *p < 0.05, **p < 0.01, ***p < 0.001 (Mann-Whitney U test). WMG, Watchmaker Genomics RRI; No, no-inhibitor control.

Across all tested SEQURNA concentrations in retinal organoids, the pattern was remarkably consistent: 0 mM DTT outperformed 5 mM which, in turn, outperformed 10 mM DTT for gene detection (Figure 2C). Importantly, DTT supplementation did not improve other RNA quality metrics-mapping efficiency remained stable (77-79% unique mapping, F=1.31, p=0.27), and splice junction detection showed no improvement (F=1.44, p=0.24). These results demonstrate that DTT, while intended to protect RNA and enzymes from oxidative damage, paradoxically impairs reverse transcription without compensatory benefits when using SEQURNA. Based on these findings, we recommend omitting DTT from the FLASH-seq protocol when using SEQURNA.

Cell type composition analysis confirmed that SEQURNA preserves biological heterogeneity comparably to commercial alternatives (Figure 2E-F). In retinal organoids, all seven major populations were identified across conditions: amacrine cells, bipolar cells, retinal ganglion cells, immature neurons, Müller glia, retinal progenitor cells, and photoreceptors. Similarly, all six major PBMC subsets were detected. Dimensionality reduction revealed minimal batch effects, with cells clustering by biological type rather than experimental condition, confirming that RNase inhibitor selection does not distort sample composition.

## Conclusions

Our systematic evaluation establishes SEQURNA as a superior alternative to commercial RRIs for FLASH-seq, with 0.75-1 U/μl providing optimal balance between transcriptome coverage and data quality. The performance advantage was particularly pronounced in PBMCs, consistent with observations that effective RNase inhibition becomes increasingly critical in low-RNA cells where transcript losses have proportionally greater impact [13]. The elevated splice junction complexity in PBMCs compared to retinal organoids likely reflects biological differences, as circulating immune cells undergo extensive alternative splicing generating thousands of unique junctions per cell [12,14].

Perhaps most strikingly, DTT supplementation at routinely used concentrations significantly impaired gene detection without improving RNA quality, challenging decades of established practice. DTT is universally included based on the rationale that it stabilizes enzyme activity and inhibits RNases by reducing critical disulfide bonds. Our data suggest competing biochemical effects may outweigh these intended benefits. Notably, this applies specifically to SEQURNA, which unlike protein-based RRIs does not require reducing agents for functionality.

The cost-effectiveness of SEQURNA is compelling at scale. At our recommended 1 U/μl concentration, SEQURNA costs approximately 21 CHF per 384-well plate compared to 70 CHF for Takara RRI, representing a 70% reduction-substantial savings for facilities processing hundreds of thousands of cells annually. Several limitations should be acknowledged: our evaluation focused exclusively on FLASH-seq, and performance may differ in platforms with distinct buffer compositions or thermal profiles. Additional validation in tissues with high endogenous RNase content (e.g., pancreas) would strengthen generalizability. Furthermore, SEQURNA’s protective range against the full RNase spectrum remains to be comprehensively profiled. Despite these limitations, our findings establish SEQURNA at 1 U/μl without DTT as an optimized formulation that simultaneously improves performance and reduces costs.

## Materials and Methods

Human PBMCs from healthy donors (Stem Cell Technologies) were thawed, stained with FITC-conjugated anti-CD45 antibody (clone HI30, BD Biosciences), and sorted on a FACSAria Fusion (100 μm nozzle) excluding dead cells (propidium iodide-positive). Human retinal organoids were dissociated using Neural Tissue Dissociation Kit P (Miltenyi Biotec) at 37°C with gentle agitation until single-cell suspension was achieved (∼20–25 min), then sorted excluding debris, doublets, and dead cells. Single cells were sorted into 384-well plates containing 1 μl lysis buffer (Triton X-100, dNTPs, SMART-dT30VN primer, RNase inhibitor at variable concentrations, betaine, dCTP, and FLASH-seq TSO). After sorting, plates were stored at −80°C until processing. For RT-PCR, plates were incubated at 72°C for 3 min, then 4 μl RT-PCR mix containing Maxima H-reverse transcriptase (ThermoFisher) and KAPA HiFi ReadyMix (Roche) was added. Reactions were performed at 50°C for 60 min followed by 21–22 PCR cycles.cDNA was purified using SeraMag beads (0.8:1 ratio), normalized to 200 pg/μl, tagmented with in-house Tn5 transposase, and sequenced on NextSeq 550 (75-8-8-75 read configuration). A detailed protocol is available on Protocols.io.

Raw sequencing data were demultiplexed using bcl-convert (v4.3.13) and aligned to the human reference genome (GRCh38) using STAR (v2.7.11b) with GENCODE v44 annotations. Each well was processed independently, with gene-level quantification and splice junction detection performed during alignment. Cells with ≥75,000 uniquely mapped reads were retained and computationally downsampled to median sequencing depth to enable fair comparison across conditions. Quality metrics including unique mapping rate, multimapping rate, mitochondrial content, and splice junction counts were extracted from STAR output files. A composite quality score was calculated as (% Uniquely Mapped) × (1 − Mismatch Rate/100) × (% Annotated Splices/100). Cell types were annotated using canonical marker gene scoring in Scanpy, with retinal organoid populations (rods, cones, Müller glia, amacrine, bipolar, retinal ganglion cells, immature neurons) and PBMC subsets (T cells, B cells, NK cells, monocytes, dendritic cells, platelets) identified based on established markers. Statistical comparisons used Mann-Whitney U tests; two-way ANOVA assessed RRI × DTT interactions. Full methods including marker gene lists and quality filtering parameters are provided in Supplementary Methods.

## Declarations

### Ethics approval and consent to participate

Human retinal organoids. Tissue samples are fully anonymized, were obtained in accordance with the tenets of the Declaration of Helsinki and all experimental protocols were approved by the local ethics committee (Ethikkommission Nordwest- und Zentralschweiz (EKNZ)). All the relevant information, including Ethics are described on: https://hpscreg.eu/cell-line/lOBi00l-A

### Consent for publication

Not applicable.

### Availability of data and materials

All data generated or analysed during this study are included in this published article [and its supplementary information files].

### Authors’ contributions

Conceptualization: SP. Data analysis: IK, SP. Performed experiments: IK, SP, MR, SC. Writing: IK, SP. All authors read and approved the final manuscript.

## Acknowledgements

We thank S. Stefanova and M. Esposito for their help in sorting human retinal organoid cells. We thank Larissa Utz from the Human Organoid Platform at IOB for providing us with the organoids used in this study.

