## Supplementary Methods for "SEQURNA enhances FLASH-seq gene detection while eliminating DTT dependence"

### Supplementary Methods for the manuscript "SEQURNA enhances FLASH-seq gene detection while eliminating DTT dependence".

**Preparation and FACS sorting of human peripheral blood mononuclear cells**

Human PBMCs from 2 healthy donors were purchased from Stem Cell Technologies (cat. # 70025.1). Prior to FACS sorting, cells were thawed, directly resuspended in 1 x PBS + 2% FBS and stained with a FITC-conjugated mouse anti-human CD45 antibody (clone HI30, BD Biosciences) for 20 min on ice. Unbounded antibodies were washed away by adding 10 ml of 1 x Roswell Park Memorial Institute 1640 medium (RPMI, ThermoFisher Scientific) + 2% FBS, centrifuged for 5 min at 300 x g, resuspended in 1 x PBS + 0.04% BSA, strained through a 40-µm filter (pluriSelect) and stained for 5 min with propidium iodide (1 mg/ml, ThermoFisher Scientific) at room temperature to label dying cells. A MoxiGo 532 instrument (Orflo Technologies) was used for counting.

Single cells were sorted on a FACSAria Fusion instrument (BD Biosciences) with a 100-µm nozzle. Only singlets that were FITC+PI- were sorted into 384-well plates (LoBind twin.tec, Eppendorf) containing 1 µl of lysis buffer.

**Preparation and FACS sorting of human retinal organoids**

Human retinal organoids were cultured as described in Hahaut et al. [11]. Organoids of the same age and same batch were pooled together, organoids of different ages were processed in different experiments and different days. For every age and experimental day, several conditions were tested and always included our reference condition ("Takara v1" RRI) and negative control (no RNAse inhibitor added to the lysis and RT-PCR mix). On the day of the experiment 5 organoids were pooled together and washed once with 1 ml Ringer solution (Carl Roth). The Neural Tissue Dissociation Kit P (Miltenyi Biotec) was used to dissociate them into single cells. After removing the Ringer solution, an enzymatic dissociation mix composed of 50 µl enzyme P and 950 µl buffer X was added. The tube was incubated on a SmartBlock thermoblock (Eppendorf) at 37°C with gentle agitation (500 rpm). The solution was mixed with a wide-bore P1000 pipette at regular intervals to facilitate cell dissociation. The reaction was stopped as soon as the organoids showed signs of dissociation (~20-25 min) by adding 15 µl Stop solution (5 µl enzyme A + 10 µl buffer Y). Samples were mixed gently by inversion and returned to the thermoblock for 10 min until the tissue appeared almost completely digested and cell aggregates deemed to be minimal. The cell suspension was briefly put on ice before being spun for 5 min at 300 x g in a pre-refrigerated (4°C) centrifuge (Eppendorf). The supernatant was carefully removed without disturbing the pellet, before performing one wash with 1 x PBS pH 7.4 + 0.04% BSA (PBS: ThermoFisher Scientific; MACS BSA: 10% solution, Miltenyi Biotec). Supernatant was removed again and cells were resuspended in 1 x PBS pH 7.4 + 0.04% BSA and sequentially strained through a 70-μm and then a 40-μm filter (pluriSelect). Viability staining was carried out with propidium iodide (1 mg/ml, ThermoFisher Scientific) at room temperature and using a 1:500 dilution, to label dying cells. Single cells were sorted on a FACSAria Fusion instrument using a 100-μm nozzle, as described above. Only debris, doublets and PI+ (i.e., dead) cells were excluded to avoid introducing any selection bias.

**Lysis buffer preparation**

Lysis buffer was dispensed in single wells with the I.DOT instrument (Dispendix), the plate sealed and stored at -20℃ until needed. The lysis buffer had different compositions, depending on which RNAse inhibitor was used for the experiment. For both Takara RRI and Watchmaker Genomics conditions, the lysis buffer had the following composition: 0.02 μl Triton X-100 (10% v/v, Sigma-Aldrich), 0.24 μl dNTP mix (25 mM of each, Roth), 0.018 μl SMART-dT30VN (5´Bio-AAGCAGTGGTATCAACGCAGAGTACT$_{30}$VN-3´; Bio = C6-linker biotin; 100 μM, IDT), 0.03 μl of the corresponding RNAse inhibitor (40 U/μl), 0.012 μl dithiothreitol (DTT, 100 mM; Sigma-Aldrich), 0.2 μl betaine (5 M, Sigma-Aldrich), 0.09 μl dCTP (100 mM, ThermoFisher Scientific), 0.092 μl FLASH-seq template-switching oligonucleotide (FS TSO, 5′ Bio-AAGCAGTGGTATCAACGCAGAGTACrGrGrG-3′, Bio = C6-linker biotin; 100 μM, IDT) and water to 1 μl final volume.

For the SEQURNA conditions, the lysis buffer had the following composition: 0.02 μl Triton X-100 (10% v/v, Sigma-Aldrich), 0.24 μl dNTP mix (25 mM of each, Roth), 0.018 μl SMART-dT30VN (5´Bio-AAGCAGTGGTATCAACGCAGAGTACT$_{30}$VN-3´; Bio = C6-linker biotin; 100 μM, IDT), variable amounts of the SEQURNA inhibitor (50 Unit mass/μl), 0.2 μl betaine (5 M, Sigma-Aldrich), 0.09 μl dCTP (100 mM, ThermoFisher Scientific), 0.092 μl FLASH-seq template-switching oligonucleotide (FS TSO, 5′ Bio-AAGCAGTGGTATCAACGCAGAGTACrGrGrG-3′, Bio = C6-linker biotin; 100 μM, IDT) and water to 1 μl final volume.

After cell sorting, plates were sealed with aluminium foil seals (VWR International) for cold temperature storage and immediately placed in a -80°C freezer until ready for further processing.

**RT-PCR reaction**

Plates were removed from the -80°C storage, immediately transferred to a pre-heated Mastercycler thermocycler (Eppendorf), incubated for 3 min at 72°C and then placed on a metal block kept in an ice bucket. Four microliters of RT-PCR mix were added using the I.DOT. The RT-PCR mix had a different composition, depending on which RNAse inhibitor was used. For both Takara RRI and Watchmaker Genomics conditions the reaction mix had the following composition:: 0.238 µl dithiothreitol (DTT, 100 mM; Sigma-Aldrich), 0.8 µl betaine (5 M, Sigma-Aldrich), 0.046 µl magnesium chloride (1 M, Ambion), 0.096 µl of the corresponding RNAse inhibitor (40 U/µl), 0.05 µl Maxima H- reverse transcriptase (200 U/µl, ThermoFisher Scientific), 2.5 µl KAPA HiFi Hot-Start ReadyMix (2 x, Roche) and nuclease-free water to 4 µl final volume. The plate content was briefly vortexed and spun down, before starting the RT-PCR reaction: 60 min at 50°C, 98°C for 3 min, then N cycles of (98°C for 20 sec, 67°C for 20 sec, 72°C for 5 min). The number of PCR cycles depends on the cell type and on the protocol. We used 21 cycles for human retinal organoids and 22 cycles for PBMC.

**Sample cleanup, QC and normalization**

Samples were cleaned up using SeraMag SpeadBeads™ (Cytiva) containing 18% w/v polyethylene glycol MW = 8000 (Sigma-Aldrich) on a Fluent 780 automated workstation (Tecan, Switzerland), using a 0.8:1 ratio of beads:cDNA  Purified cDNA was eluted in 15 µl nuclease-free water.

Sample cDNA concentration was measured using the Quant-iT™ PicoGreen™ dsDNA assay kit (ThermoFisher Scientific) on black Nunc™ F96 MicroWell™ polystyrene plates (ThermoFisher Scientific), according to the manufacturer's instructions, except that we used half of the recommended reaction volumes and a 1:800 dilution of the fluorescent dye, to reduce costs. Fluorescence measurements were recorded on a Hidex Sense instrument (Hidex).

The concentration returned by the PicoGreen™ measurement was used as an input for normalizing each sample to a final concentration of 200 pg/µl using the Fluent liquid handling robot and the I.DOT dispenser. This plate represented our working dilution cDNA plate, used for library preparation.

**Library preparation**

Tn5 transposase produced by the EPFL Protein Facility (Lausanne) was used in all experiments. The tagmentation reaction mix contained 1 µl cDNA (200 pg/µl), 0.8 µl 100% dimethylformamide (DMF, Sigma-Aldrich), 0.8 µl 5 x TAPS buffer (50 mM TAPS-NaOH pH 7.3 at 25°C, 25 mM MgCl2; 0.22 µm filtered), 0.3 µl Tn5 transposase (~2 µM) and nuclease-free water to 4 µl final volume. After adding all the components, the plate was gently vortexed, spun down and incubated for 8 min at 55°C. The Tn5 was inactivated with 1 µl 0.2% SDS, followed by a 5 min incubation at room temperature. The indexing PCR mix was then added to the tagmented cDNA: 0.2 µl KAPA HiFi DNA Polymerase (1 U/µl), 0.3 µl dNTP mix (10 mM), 2 µl KAPA HiFi Buffer (5 x) (all part of the KAPA HiFi PCR kit, Roche), 2 µl pre-mixed N7xx and S5xx index adaptors (5 µM each) and water to 5 µl final volume. The enrichment PCR reaction was carried out as follows: 72°C for 3 min, 95°C for 30 sec, then 12 cycles of (95°C for 10 sec, 55°C for 30 sec, 72°C for 30 sec), 72°C for 5 min, 4°C hold.

**Sequencing**

After PCR, cells were pooled together in a 1.5-ml LoBind tube (Eppendorf) and cleaned up with a 0.8:1 SeraMag SpeadBeads™ containing 18% w/v polyethylene glycol (MW = 8000). Elution was performed in nuclease-free water and 1 µl was used for measuring the concentration with the Qubit™ dsDNA high sensitivity assay kit (ThermoFisher Scientific), while the size was assessed with a high sensitivity DNA chip on a 2100 Bioanalyzer instrument.

Libraries prepared with different sets of index primers were pooled together and sequenced using a NextSeq™ 550 High Output kit v2.5 (150 cycles) with read mode 75-8-8-75.

**Demultiplexing and FASTQ generation**

Raw sequencing data (BCL files) were demultiplexed and converted to FASTQ format using bcl-convert (v4.3.13, Illumina) with default quality filtering. Lane splitting was disabled to combine reads across lanes, and FASTQ files were compressed using gzip compression level 1 for efficient storage.

**Sequence alignment and gene quantification**

Paired-end reads were aligned to the human reference genome (GRCh38, Homo_sapiens.GRCh38.dna.primary_assembly.fa) using STAR (v2.7.11b) with a pre-built genome index generated from GENCODE v44 annotations (gencode.v44.annotation.gtf). Each well of the 384-well plate was processed independently as an individual sample, consistent with the plate-based single-cell RNA-seq design of FLASH-seq. To optimize computational efficiency, the genome index was loaded into shared memory and maintained across all samples within a plate. Alignment was performed with the following parameters: output as coordinate-sorted BAM files (`--outSAMtype BAM SortedByCoordinate`), gene-level quantification enabled (`--quantMode GeneCounts`), 8 threads per sample (`--runThreadN 8`), and BAM sorting RAM limited to 10 GB. Gene-level read counts were extracted from STAR's ReadsPerGene.out.tab output files using unstranded counts. Splice junctions were obtained from SJ.out.tab files, with high-confidence junctions defined as those supported by ≥2 uniquely mapped reads.

**Cell filtering and quality control**

Wells were considered to contain successfully captured cells if they yielded ≥75,000 uniquely mapped reads; wells below this threshold were excluded from downstream analysis. Mapping quality metrics were extracted from STAR Log.final.out files, including the percentage of uniquely mapped reads, multimapped reads, reads unmapped due to short length, and mismatch rate per base. A composite quality score was calculated as:

*Quality Score = (% Uniquely Mapped) × (1 − Mismatch Rate/100) × (% Annotated Splices/100)*

A degradation index was computed as the sum of the percentage of unmapped short reads and the percentage of novel splice junctions, where elevated values indicate potential RNA degradation artifacts.

Mitochondrial transcript content was calculated as the percentage of reads mapping to mitochondrial genes (MT- prefix). Ribosomal RNA content was determined from reads mapping to genes with RPS, RPL, MRPS, or MRPL prefixes. Ensembl gene identifiers were converted to gene symbols using the MyGene.info API.

**Depth normalization**

To enable fair comparison across conditions with varying sequencing depths, all samples passing the initial read threshold were computationally downsampled to the median sequencing depth across all cells. Downsampling was performed by random sampling without replacement from the pool of mapped reads, preserving the original distribution of reads across genes while equalizing total counts.

##### Outlier detection and additional quality filtering

Outlier cells were identified using the Median Absolute Deviation (MAD) method applied to gene detection counts within each plate and RRI condition group. Cells with modified Z-scores exceeding 3.5 MADs from the median were flagged as outliers. A minimum of 5 cells per group was required for outlier detection. Additionally, cells with fewer than 500 detected genes (300 for DTT-specific analyses) were excluded as low-quality.

##### Dimensionality reduction and visualization

Single-cell analysis was performed using Scanpy (v1.9.x) in Python. Gene expression matrices were normalized to 10,000 counts per cell and log-transformed. Highly variable genes (n=3,000) were identified using the Seurat v3 method with batch correction for cell source (retinal organoid vs. PBMC) to prevent tissue-specific bias. Following scaling (maximum value capped at 10), principal component analysis was performed retaining 50 components. A k-nearest neighbor graph was constructed using 12 neighbors and 30 principal components, followed by UMAP embedding (min_dist=0.3, spread=1.0) for visualization.

##### Cell type annotation

Cell types were annotated using marker gene scoring. For retinal organoids, canonical markers were used for rods (RHO, GNAT1, PDE6B, NRL, ROM1), cones (OPN1MW, OPN1LW, OPN1SW, ARR3, GNAT2), Müller glia (RLBP1, GLUL, VIM, AQP4, CRYAB), amacrine cells (GAD1, GAD2, TFAP2A, PAX6, SLC6A9), bipolar cells (VSX1, VSX2, PRKCA, GRM6, TRPM1), retinal ganglion cells (POU4F1, RBPMS, SNCG, THY1, NEFL), and immature neurons (DCX, NEUROD1, STMN2, TUBB3, ELAVL3). For PBMCs, markers included T cells (CD3D, CD3E, TRAC, IL7R), B cells (MS4A1, CD79A, CD79B, BANK1), NK cells (NKG7, GNLY, KLRD1, PRF1), monocytes (LYZ, S100A8, S100A9, LST1, VCAN), dendritic cells (FCER1A, CST3, LILRA4), and platelets (PPBP, PF4, NRGN). Each cell was assigned to the cell type with the highest marker gene score using Scanpy's `score_genes` function.

##### Statistical analysis

Comparisons between RRI conditions were performed using the Mann-Whitney U test. Effect sizes were calculated as rank-biserial correlation coefficients ($r = 1 - 2U/n_1n_2$). For analyses involving multiple factors (RRI concentration × DTT concentration), two-way ANOVA was used to assess main effects and interactions. Statistical significance was defined as $p < 0.05$, with significance levels denoted as: * $p < 0.05$, ** $p < 0.01$, *** $p < 0.001$.
