## Supplementary Figures for "SEQURNA enhances FLASH-seq gene detection while eliminating DTT dependence"

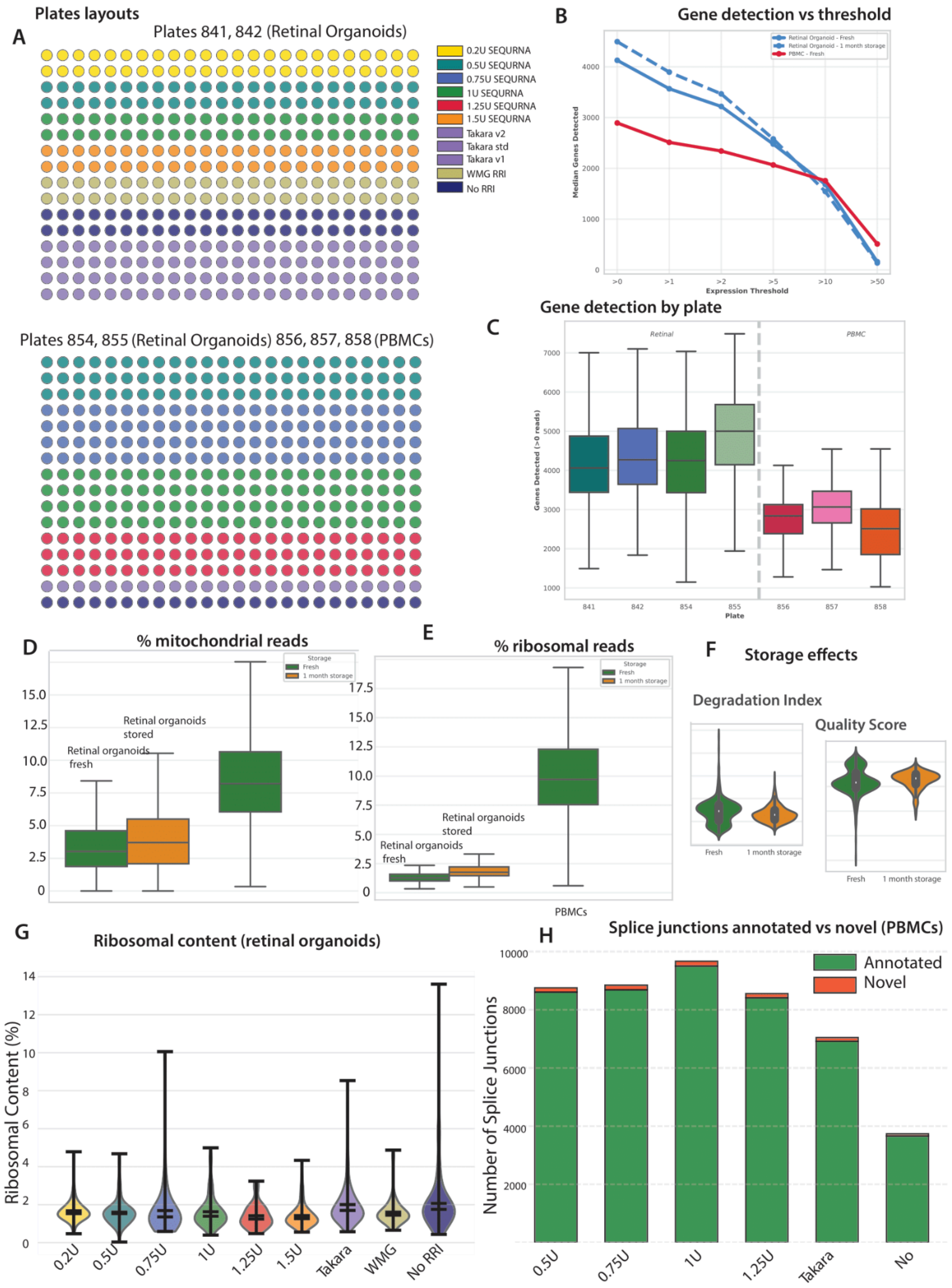

A
Plates layouts
Plates 841, 842 (Retinal Organoids)
0.2U SEQURNA
0.5U SEQURNA
0.75U SEQURNA
1U SEQURNA
1.25U SEQURNA
1.5U SEQURNA
Takara v2
Takara std
Takara v1
WMG RRI
No RRI
Plates 854, 855 (Retinal Organoids) 856, 857, 858 (PBMCs)
B
Gene detection vs threshold
Retinal Organoid - Fresh
Retinal Organoid - 1 month storage
PBMC - Fresh
Median Genes Detected
Expression Threshold
C
Gene detection by plate
Retinal
PBMC
Genes Detected (>0 reads)
Plate
D
% mitochondrial reads
Storage
Fresh
1 month storage
Retinal organoids stored
Retinal organoids fresh
E
% ribosomal reads
Retinal organoids stored
Retinal organoids fresh
PBMCs
F
Storage effects
Degradation Index
Quality Score
Fresh
1 month storage
G
Ribosomal content (retinal organoids)
Ribosomal Content (%)
0.2U
0.5U
0.75U
1U
1.25U
1.5U
Takara
WMG
No RRI
H
Splice junctions annotated vs novel (PBMCs)
Annotated
Novel
Number of Splice Junctions
0.5U
0.75U
1U
1.25U
Takara
No

**Supplementary Figure 1. Experimental design, storage stability, and additional quality metrics. (A)** Plate layouts showing experimental conditions across 384-well plates. Plates 841 and 842: retinal organoids with SEQURNA concentrations (0.2–1.5 U), commercial RRIs (Takara v1, Takara v2, Takara std, WMG), and no-RRI controls. Plates 854–855: retinal organoids; plates 856–858: PBMCs. **(B)** Median genes detected at increasing expression thresholds comparing fresh retinal organoids, stored retinal organoids (1 month at −80°C), and fresh PBMCs. **(C)** Gene detection by plate for retinal organoids (plates 841, 842, 854, 855) and PBMCs (plates 856, 857, 858). **(D)** Mitochondrial transcript content (%) comparing fresh versus 1-month stored retinal organoid samples. **(E)** Ribosomal transcript content (%) for fresh and stored retinal organoids and PBMCs. **(F)** Storage effects on data quality: degradation index (left) and composite quality score (right) comparing fresh versus 1-month stored samples. **(G)** Ribosomal content distribution across RRI conditions in retinal organoids. Violin plots show distribution with median indicated. **(H)** Splice junction classification in PBMCs showing annotated versus novel junctions across conditions. The high ratio of annotated junctions (>98%) indicates minimal spurious splicing artifacts.

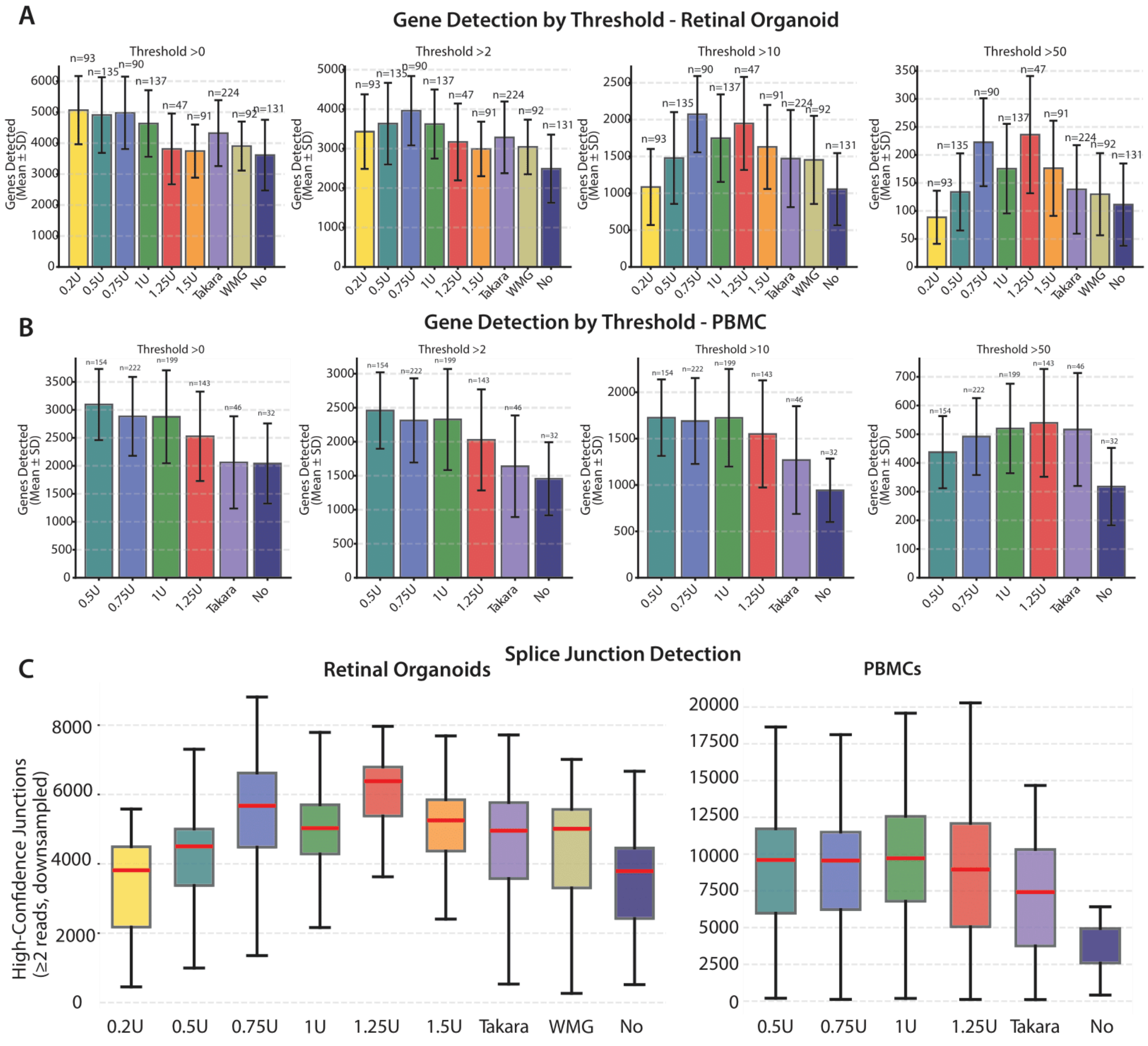


**Supplementary Figure 2. Gene detection across expression thresholds and splice junction detection. (A)** Gene detection (mean ± SD) in retinal organoids across SEQURNA concentrations and controls at four expression thresholds (>0, >2, >10, >50 reads). Sample sizes indicated above bars. At higher thresholds (>10, >50 reads), mid-to-high SEQURNA concentrations (0.75–1.25 U) demonstrate superior performance, indicating reduced low-level background noise while preserving robustly expressed transcripts. **(B)** Gene detection (mean ± SD) in PBMCs across conditions and expression thresholds. **(C)** High-confidence splice junction detection (≥2 supporting reads, downsampled) in retinal organoids (left) and PBMCs (right). PBMCs exhibit 1.7–2.0-fold higher junction counts than retinal organoids across all conditions, likely reflecting extensive alternative splicing and V(D)J recombination in immune cells. Boxplots show median (red line), interquartile range, and outliers. WMG, Watchmaker Genomics RRI; No, no-inhibitor control.
